# Networked information interactions of epileptic EEG based on symbolic transfer entropy

**DOI:** 10.1101/543496

**Authors:** Wenpo Yao, Jun Wang

## Abstract

Identifying networked information exchanges among brain regions is important for understanding the brain structure. We employ symbolic transfer entropy to facilitate the construction of networked information interactions for EEGs of 22 epileptics and 22 healthy subjects. The epileptic patients during seizure-free interval have lower information transfer in each individual and whole brain regions than the healthy subjects. Among all of the brain regions, the information flows out of and into the brain area of O1 of the epileptic EEGs are significantly lower than those of the healthy (p<0.0005), and the information flow from F7 to F8 (p<0.00001) is particularly promising to discriminate the two groups of EEGs. Moreover, Shannon entropy of probability distributions of information exchanges suggests that the healthy EEGs have higher complexity and irregularity than the epileptic brain electrical activities. By characterizing the brain networked information interactions, our findings highlight the long-term reduced information exchanges, degree of brain interactivities and informational complexity of the epileptic EEG.

## 1. Introduction

The emerging field of network science, derived or evolved from the physical real world, contributes to revealing the underlying endogenous structural information, like social or economic networks [1, 2], networked physiology [3, 4] and so on. Human brain, a collection of purportedly 100 billion neurons and 10× more glial cells [5], is arguably a dynamical complex network characterized with formidably complicated structure, interactive causalities, and brain behaviors are subject to various internal and external factors like stress, pathological conditions and so on. Modern brain imaging techniques, recording magnetic or electrical brain activities, produce increasingly large anatomical or functional connected datasets. The networked technique therefore have been gained particularly popularity in elucidating the brain dynamical complex structures [6, 7] that are featured by like small-world topology or highly connected hubs at the whole-brain scale or at a cellular scale [8]. Brain networks have underscored value for identifications of neurological and psychiatric alterations, and understanding the evolving human brain networks contributes the elucidation of physiological and pathological mechanisms, such as epilepsy [9], depression [10], schizophrenia [11] etc.

To identify and quantify interaction or connectivity among brain regions are neurobiologically meaningful and challenging for understanding the structural dynamics of brain [12]. Quantitative causality is a fundamental issue for understanding dynamical information and tapping into underlying structure of networks [13]. To tackle the existence of functional or effective connectivity between two or among several brain regions, several methods including correlation, cross-correlation, Granger causality, phase slope index [14], or information-theoretic approaches [15] are proposed to characterize synchronization, coupling or interactive relationships. Among these analytical tools, transfer entropy proposed by T. Schreiber [16] is derived to quantify statistical coherence or interaction between (sub)systems, and the the asymmetric parameter effectively distinguish driving and responding factors. Since then, the transfer entropy has been gaining growing popularity in several areas, particularly the neurological science, for quantifying the information exchanges or understanding the causal relationships or effective connectivity across data modalities like EEG [17, 18], MEG [19, 20], fMRI, and so on [21, 22].

The epileptic process can be regarded as variable-scale networked phenomenon, which is supported by growing reports [9]. To construct networked information connectivity for epileptic EEGs, we employ symbolic transfer entropy to identify informational exchanges between different brain regions, and we measure the complexity of information exchanges based on entropy method.

## 2. Methodology

### 2.1. Transfer entropy

Transfer entropy naturally incorporates directional statistical coherence between two (sub)systems, it does not assume any particular model for systems, this advantage is especially relevant in coherence detection for nonlinear interactions under unknown structural information [19].

For evolving time series *X* and *Y*, approximated by Markov processes, information entropy of *X* is 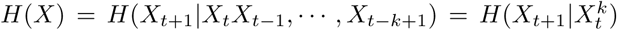. The transfer entropy from *Y* to *X* is essentially the declined information of *H*(*X*) leaded by *Y* as 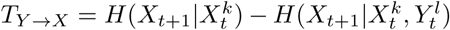 where transition probabilities are expressed by joint probabilities in experimental time series analysis, and it is mathematically expand to Eq. 2 where *l* could be *l* = *k* or *l* = 1 for most natural choices [16].

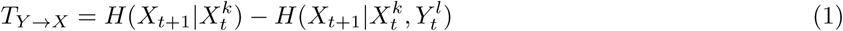

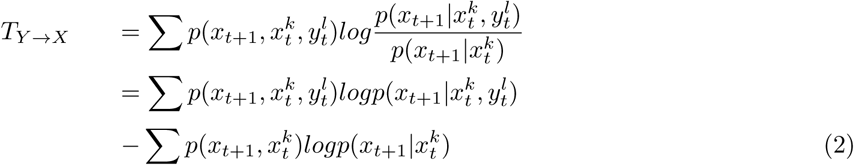

Transfer entropy, incorporating directional and dynamical information, is inherently asymmetric in mathematics, *T*_*Y* → *X*_ ≠*T*_*X*→*Y*_, and hence could be used to determine driving or responding factor by identifying direction of information flow.

The non-parametric method does not require priori assumptions of whether linear or nonlinear dependence although it is mathematically equivalent to Granger Causality [23, 24]. However, the estimation of transition probabilities from the data is not trivial, and solutions for this issue include entropy estimators [25], like the step kernel estimation which is employed by T. Schreiber [16] in the original paper. Another alternative solution is symbolic time series analysis [26] that transforms raw time series with continuous distributions into symbolic sequence containing discretized symbols from some alphabet [17, 27] to simplify the calculation of probability distributions. The symbolic transfer entropy refers to deviation of the cross-Markovian properties of symbolic sequences of ordinal patterns instead of the raw time series.

### 2.2. Transfer entropy based on permutation

In the permutation-based symbolic method, Staniek, M. and K. Lehnertz [17] combine transfer entropy and symbolic permutation to simplify the calculation of transition probabilities and reduce high demands on the data and improve insensitiveness to noise contributions. The permutation method, originally proposed in permutation entropy [28, 29], is to map time series into ordinal patterns, and it inherits the temporal structural information of system dynamics and directly applies to arbitrary real-world data.

Given time series *X*(*i*), multi-dimension phase space is constructed as 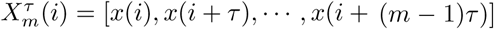 for dimension *m* and delay *τ*, elements in each vector are reordered according to their relative values *x*_*mτ*_ (*j*_1_) ≤ *x*_*mτ*_ (*j*_2_) ≤ ⋯ ≤ *x*_*mτ*_ (*j*_*i*_). The indexes of original elements constitute order patterns *π*_*j*_ = {*j*_1_, *j*_2_, ⋯, *j*_*i*_} whose upper bound is *m*!, and permutations of *m*=2 and 3 are illustrated in Fig. 1. This order pattern scheme, inheriting the causal information without any further model assumptions, is widely adopted to simplify time series analysis [29, 30, 31].

**Figure 1:**
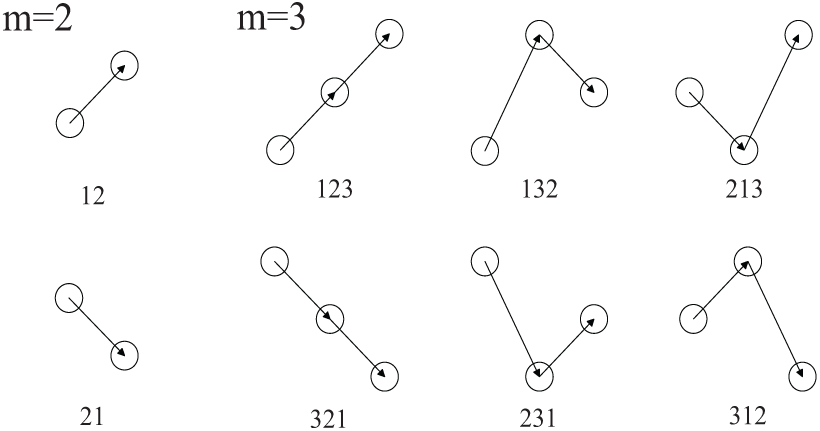
Order patterns when *m*=2 and 3.

## 3. Results

22 epilepsy patients (aged from 14 to 51 years, 30.0±13.1 years) and 22 healthy volunteers (aged from 15 to 49 years, 26.9±8.9 years) from Nanjing General Hospital of Nanjing Military Command were recruited for EEG collection [27, 30, 32]. Following the standard 10-20 system, we locate 16 scalp electrodes, namely Fp1, Fp2, F3, F4, C3, C4, P3, P4, O1, O2, F7, F8, T3, T4, T5 and T6. The time duration of data recording is 1 minute and sampling frequency is 512 Hz. Eye movement artifacts and noise have been removed. All of the volunteers are in idle states during EEG collection, and the epileptic patients are all in their seizure-free intervals. Exemplary epileptic and healthy EEGs and the order patterns are illustrated in Fig. 2.

**Figure 2:**
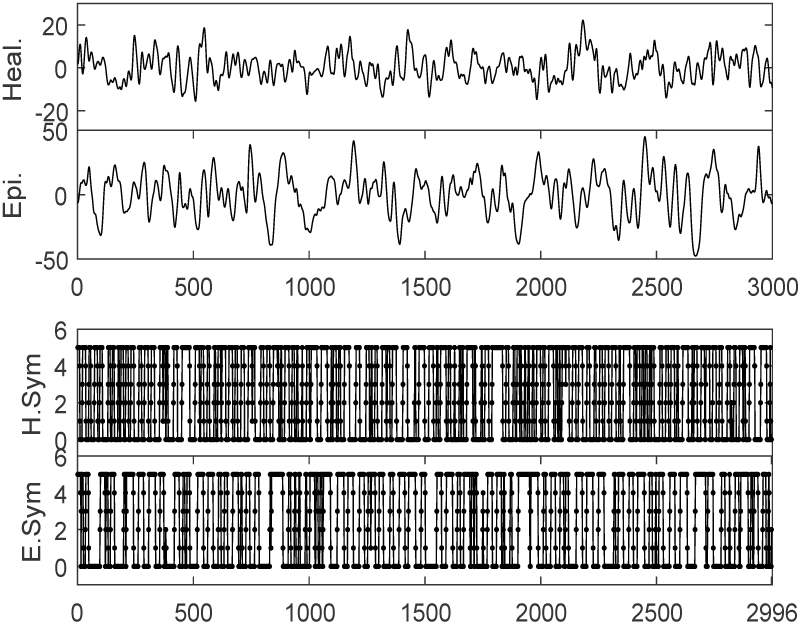
Exemplary epileptic and healthy EEGs (brain region of C3) and their order patterns.

To detect the information exchanges or causal links between different brain areas contributes to explore the brain modulation mechanism. We use the symbolic transfer entropy to construct networked information flows for epileptic brain electrical activities and characterize the effects of epilepsy on networked information interactions. Brain regions are assumed to represent the nodes of network and the information exchanges between each brain area constitute the edges of functional network. Permutation with embedding dimension of 2, 3 and 4 and delay of from 1 to 5 are adopted for symbolic transfer entropy. Exemplary directional information exchanges (*m*=3, *τ*=4) from all the brain areas are calculated to map the informational interactions between each brain region, displayed in Fig. 3.

**Figure 3:**
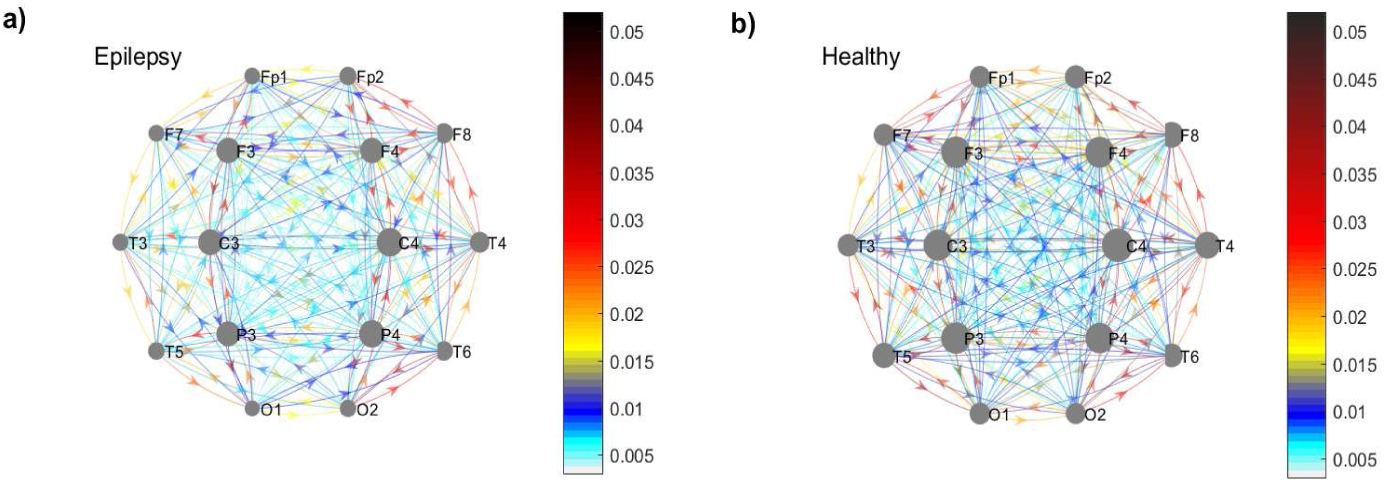
Weighted directed networks of information exchanges. a) Epileptic patients. b) Healthy subjects.

The epileptic EEGs have less information exchanges between each brain region and lower information flows in all brain areas (represented by the marker size), shown by Fig. 3. To characterize the networks of information transfer, information flows in and out of each individual brain regions are mapped to find the differences of brain informational activities between the healthy and epileptics. And we sum all the information flows as the overall degree of brain complex network, representing the degree of brain electrical activities.

Information exchanges among brain regions of the two groups of EEGs and the statistical t test for the discrimination are mapped in Fig. 4.

**Figure 4:**
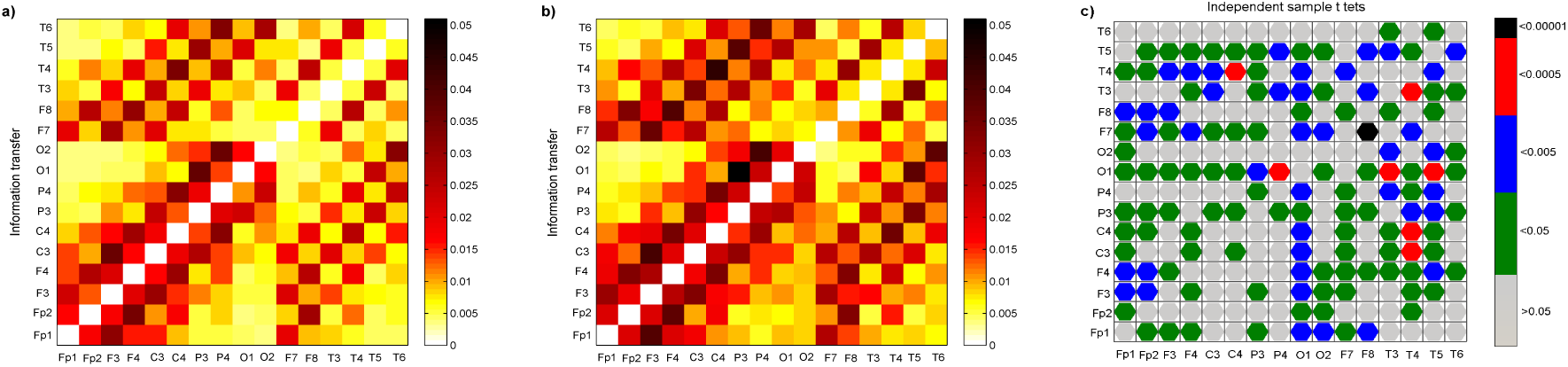
Fields of information flows between any two brain regions. a) Epileptic patients. b) Healthy subjects. c) Independent sample t tests. Information transfer is defined as *T*_*x→y*_, that information flows from brain regions in x-label to those in y-label.

Information flows between each two brain regions are asymmetric, and information exchanges of the healthy EEG are higher than those of the epileptics. According to the independent sample t tests for the discrimination of information transfer of the two groups of EEGs, shown in Fig. 4c, information transfer in gray cannot separate the two groups of EEGs (p>0.05), while there are acceptable discrimination between the healthy and epileptic EEGs in other colors (p<0.05), among which information flows in blue (p<0.005) and red (p<0.0005) are better. According to Fig. 4c, information flows out of and into the brain region of O1 have more acceptable differences, and the discrimination of information transfer from F8 to F7 is the best (p<0.00001).

Next, we calculate the information flows into and out of each region to measure the brain activities in each individual region. Taking Fp1 as an example, information flow in is 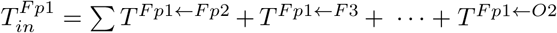, and information flows out is 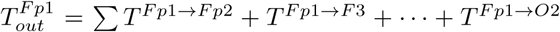. Since the information flows defined by transfer entropy between different brain regions are asymmetric, *T*_*in*_ and *T*_*out*_ are generally different. Information flows (*m*=3, *τ* =4) into and out of the 16 brain regions are illustrated in Fig. 5.

**Figure 5:**
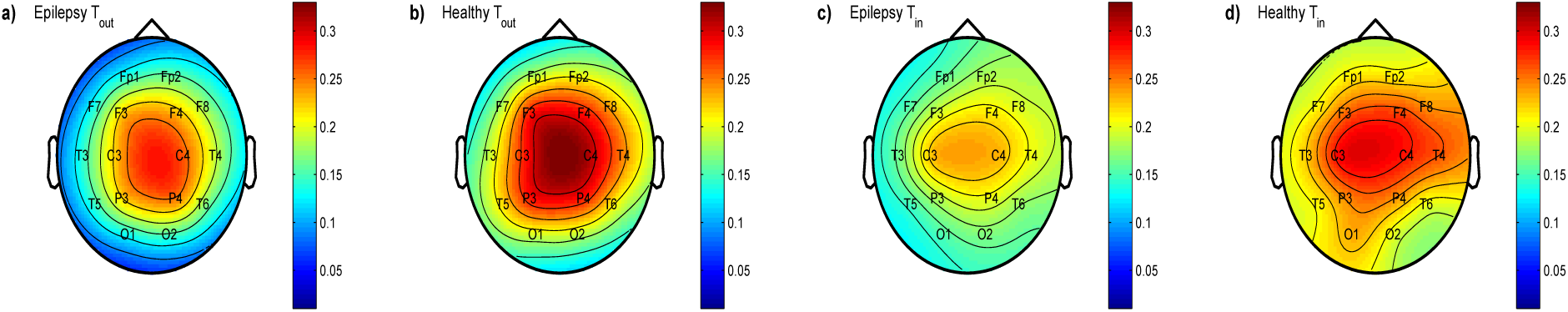
Information flows into and out of each brain region of the epileptic and healthy subjects. a) Epilepsy *T*_*out*_ b) Healthy *T*_*out*_ c) Epilepsy *T*_*in*_ d) Healthy *T*_*in*_.

Both *T*_*in*_ and *T*_*out*_ of all the individual brain regions suggest the healthy volunteers have higher degree of brain electrical activities than their epileptic counterparts, and brain areas around the central lobes have higher information interactions. However, neither information flows in and information flows out of the 16 brain regions could effectively discriminate the two kinds of EEGs (p>0.05).

We sum information exchanges of each individual brain region, taking the region of Fp1 as example 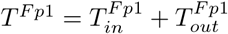, and display them in Fig. 6.

**Figure 6:**
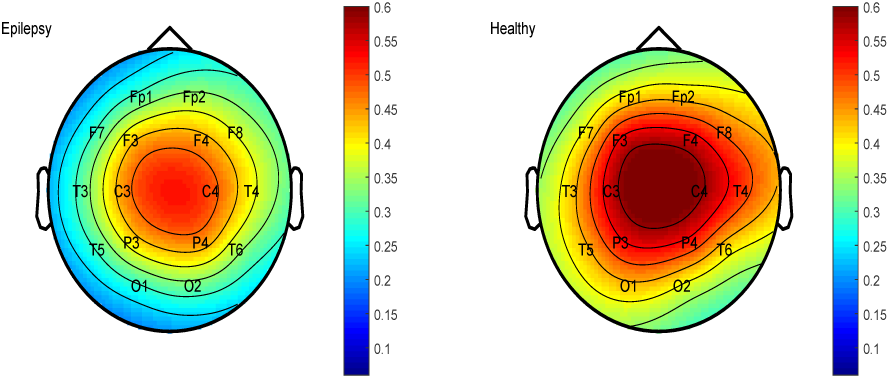
Information flows of the epileptic and healthy subjects. a) Epilepsy b) Healthy.

Informational exchanges *sum*_−_*T* = Σ*T*_*in*_ = Σ *T*_*out*_ of the whole brain regions, reflecting the degree of overall brain connective activities, are shown in Fig. 5.

**Figure 7:**
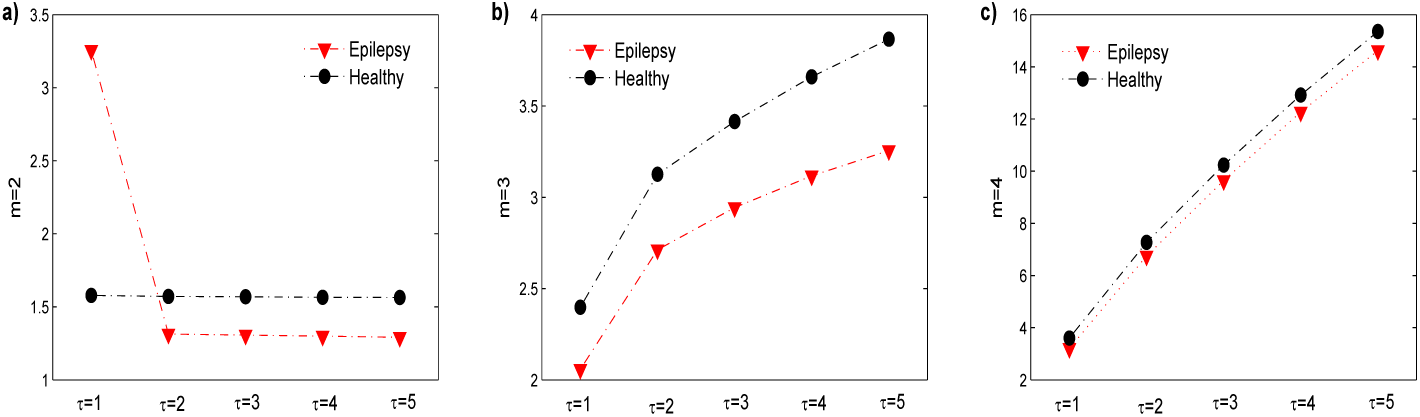
Information exchanges of the whole brain regions.

From whole brain areas, the information interactions of the epileptic EEGs are also lower than the control group, further suggesting the decreased interactivities among the brain areas of the epileptic patients. However, the discriminations between the two groups of EEGs in the whole information exchanges are not statistically acceptable (p>0.05).

Entropy measures [25], like the Shannon or Renyi entropy for static complexity and conditional entropy for dynamical complexity, are widely used to quantify the complexity of dynamical systems [33, 34]. In our contribution, we calculate the Shannon entropy for the probability of information flow from each brain region to characterize the complexity of the networked information exchanges from information-theoretic perspective. We calculate the probability of information flows of each individual brain region, like *p*_*in*_ = *T*_*in*_*/sum*_*T*_, and employ Shannon entropy for the probabilities as *H* = − Σ*p*_*in*_*logp*_*in*_, to quantify the complexity of informational interactions of EEGs, listed in Tab. 1.

**Table 1:** Complexity of informational interactions. *E*_−_*m*2 represents the complexity of epileptic EEG when *m*=2, and *H*_−_*m*2 denotes the complexity of healthy EEG when *m*=2.

| $\tau$ | 1 | 2 | 3 | 4 | 5 |
| --- | --- | --- | --- | --- | --- |
| $E\_m2$ | **3.9680** | 3.9021 | 3.9032 | 3.9037 | 3.9055 |
| $H\_m2$ | **3.9308** | 3.9314 | 3.9321 | 3.9330 | 3.9345 |
| $E\_m3$ | 3.9435 | 3.9622 | 3.9646 | 3.9664 | 3.9680 |
| $H\_m3$ | 3.9612 | 3.9746 | 3.9768 | 3.9784 | 3.9795 |
| $E\_m4$ | 3.9736 | 3.9935 | 3.9957 | 3.9964 | 3.9970 |
| $H\_m4$ | 3.9808 | 3.9950 | 3.9967 | 3.9971 | 3.9975 |

From the Tab. 1, we learn that the healthy brains have higher informational complexity than the epileptic brains. When *m*=2 and *τ*=1, the result is different from other, sharing the inconsistent results of information flows in Fig.4. The lower complexity of epileptic EEGs indicates the damage of epilepsy to the brain, which is not affected by the dimension and delay.

Our findings suggest the decreased brain interactive behaviors and reduced the complexity of brain information exchanges. Hyper-synchronous neuronal firings develop suddenly in the cerebral cortex during hallmark of recurrent seizures [35]. Accompanying epileptic seizures, there is an impairment or loss of consciousness, psychics or motor phenomena, and body vigorous shaking together with dramatic abnormal brain activities can result in physical and mental injuries. The neurological disease of epilepsy have long-term effects to the brain dynamical interactions based on our findings although the epileptics during seizure-free intervals have no significant difference in daily behaviors and brain activities. In the related reports of the present authors [30, 32, 27], the epileptic EEGs have lower nonlinearity from low-dimension dynamics, which is further verified by our networked analysis. In the nonlinearity of time irreversibility analysis [30], epileptic brain activities showed lower nonlinear dynamics especially in the brain areas of parietal lobes, which is close to the most distinct occipital lobe of O1 in our informational exchanges.

In the epileptogenic zones, one can find some high amplitude patterns in the brain electric activities during seizures. R.G. Andrzejak et al. [36] focus on nonlinear dynamical properties of epileptic brain electric activities under different pathological states, and they identify strongest nonlinear dynamics for seizure activities while suggest the nonlinearity of seizure-free EEGs to be in-between the healthy and seizure EEGs. Other followers [37, 38] adopting these groups of continuous EEGs share the outcomes that the epileptic seizure-free EEGs nonlinearity is higher than the healthy volunteers. Our findings, however, challenge the reports and suggest the healthy subjects should have more dynamical activities than the epileptics in their seizure-free intervals. A tentative explanation to the different findings could be circadian and circaseptan rhythms in human epilepsy [30, 39] that abnormally firing nerves during seizures lead to long-term damage to the brain and the epileptic seizure-free brain activities (about 20 to 30 days from the latest seizures) decrease interactions or synchronization of the brain networks, which reduce information exchanges between different brain regions and the degree of brain activities.

## 4. Conclusions

We construct networked information interactions using permutation transfer entropy for understanding the structural dynamics of brain information interactions and elucidating the underlying mechanism of the epileptic EEGs. The epileptic seizure-free EEGs have lower informational exchanges of individual and whole brain regions as well as lower information-theoretic complexity than the healthy, suggesting the long-term negative effects of epilepsy on the brain information interactions.

## 5. Acknowledgment

The project is supported by the National Natural Science Foundation of China (Grant Nos. 31671006,61771251), Jiangsu Provincial Key R&D Program (Social Development) (Grant No.BE2015700,BE2016773), Natural Science Research Major Program in Universities of Jiangsu Province (Grant No.16KJA310002).

